# A new cell culture resource for investigations of reptilian gene function

**DOI:** 10.1101/2023.09.25.559349

**Authors:** Sukhada P. Samudra, Sungdae Park, Elizabeth A. Esser, Tryggvi P. McDonald, Arianna M. Borges, Jonathan Eggenschwiler, Douglas B. Menke

**Affiliations:** Department of Genetics, University of Georgia, Athens, GA 30602, USA

## Abstract

The recent establishment of CRISPR/Cas9 gene editing in *A. sagrei* lizards makes it a powerful model system for studies of reptilian gene function. To enhance the versatility of this model, we developed an immortalized lizard fibroblast cell line (ASEC-1) for the exploration of reptilian gene function in cellular processes. We demonstrate the use of this *in vitro* system by scrutinizing the role of primary cilia in lizard Hedgehog (Hh) signaling. Through CRISPR/Cas9 mutagenesis we disrupted the *ift88* gene, which is required for ciliogenesis in diverse organisms. We find that the loss of *itf88* from lizard cells results in an absence of primary cilia, a partial derepression of *gli1* transcription, and an inability of the cells to respond to the Smoothened agonist, SAG. Through a cross-species analysis of SAG-induced transcriptional responses in cultured limb bud cells, we further determined that ∼46% of genes induced as a response to Hh pathway activation in *A. sagrei,* are also SAG-responsive in *M. musculus* limb bud cells. Our results highlight conserved and diverged aspects of Hh signaling in anoles and establish a new resource for investigations of reptilian gene function.

## Introduction

Hedgehog (Hh) proteins act as secreted ligands and regulate a large number of developmental processes in animals (McMahon et.al, 2003). Genetic studies in *Drosophila*, mice, and zebrafish have revealed that core components of the Hh signaling pathway are highly conserved between vertebrates and arthropods with some key differences. For instance, genes encoding certain components of the Hh pathway have been duplicated and have functionally diversified in vertebrates (Varjosalo & Taipale, 2008), and the contribution of some Hh pathway components to signaling has changed over evolutionary time (Chen et al., 2009; Gigante & Caspary, 2020; Huangfu et al., 2003; Wilson et al., 2009). However, detailed investigations of Hh signaling mechanisms have only been carried out in a handful of model systems. Therefore, our knowledge of Hh signaling differences among species remains limited.

Functional investigations of the Gli transcription factors, which are downstream mediators of Hh signal transduction, have demonstrated that the relative roles of Glis in mediating Hh responses differ among vertebrate species. Glis are typically regulated by the pathway to adopt either transcriptional repressor (Gli^Rep^) or activator (Gli^Act^) states. The ratio of Gli^Act^ to Gli^Rep^ in a cell determines the magnitude of its response to varying levels of Hh signals. Gli2 and Gli3 can act as either activators or repressors, depending on post-translational processing, whereas Gli1 lacks the site for proteolytic cleavage and, as a result, can only act as an activator (Falkenstein & Vokes, 2014). Within the vertebrate lineage, however, the relative roles of Gli proteins have diverged. In mice, Gli2 is the major transcriptional activator of the pathway while Gli1 plays a very minor role. Mouse Gli3 acts predominantly in transcriptional repression, although it can also function as an activator (Bai et al., 2002; Ding et al., 1998; Mo et al., 1997). In zebrafish, Gli1 is the major activator of the pathway, and Gli2 has a minor role mostly limited to repression (Huang & Schier, 2009; Karlstrom et al., 2003; Tyurina et al., 2005). Zebrafish Gli3, like its mouse homolog, functions as either an activator or a repressor, depending on developmental context (Tyurina et al., 2005). In chickens, GLI2 and GLI3 appear to act as both activators and repressors although these roles have yet to be investigated through genetic disruption (Lei et al., 2004; Persson et al., 2002; Stamataki et al., 2005).

Another distinct and unique feature of vertebrate Hh signaling is the requirement of primary cilia for the transduction of the pathway (Ben et al., 2011; Huang & Schier, 2009; Huangfu et al., 2003; Park et al., 2006; Yin et al., 2009). In vertebrates, these microtubule-based cellular appendages act as specialized centers for Hh signal transduction. Primary cilia mediate Hh signaling by modulating Gli activator and repressor activities, and proper production of the transcriptional activator and repressor forms of Gli2 and Gli3 require cilia (Eggenschwiler & Anderson, 2007; Santos & Reiter, 2014). In mice, primary cilia play a central role in ligand-induced Hh pathway activation and a relatively smaller role in repression of the pathway in the absence of ligand stimulation (Huangfu et al., 2003; Huangfu & Anderson, 2005; Ocbina et al., 2009; Wang et al., 2019). In contrast, it appears the more important role of cilia in zebrafish Hh signaling is in prevention of inappropriate pathway activation by restricting the activity of the Gli1 activator (Karlstrom et al., 2003). Zebrafish also require cilia to activate pathway targets that require moderate-to-high levels of Hh signaling (Huang & Schier, 2009). Chicken *Talpid3* mutants fail to develop primary cilia (Lewis et al., 1999) and their molecular phenotypes suggest that in chickens, as in mice, cilia play a major role in activation of the Hh pathway and a less important role in Hh pathway repression.

Some aspects of Hh signal transduction appear to be changing over vertebrate evolution, but detailed studies of Hh signaling have only been performed in a few model species, preventing a deeper understanding of when and how changes in Hh signaling mechanisms have evolved in vertebrates. One major group of amniotes that remains largely unstudied are squamate reptiles, the group that comprises lizards and snakes. This group diverged from mammals ∼320 million years ago and from avians ∼290 million year ago (Hedges et al., 2006). Thus, squamates are distantly related to commonly studied amniotes, including mouse and chick. Experimental manipulation of Hh in squamate reptiles has revealed roles of Hh signaling in tooth morphogenesis, cartilage development, lung development, and lizard tail regeneration (Handrigan & Richman, 2010; Lozito et al., 2021; Lozito & Tuan, 2015, 2016; Palmer et al., 2021; Vonk et al., 2022). In addition, evolutionary reduction in limb length and limb loss in snakes has been associated with the progressive loss of function of a limb specific cis-regulatory element of the *Shh* gene (Kvon et al., 2016; Leal & Cohn, 2016). These studies highlight the importance of Hh in reptile development and evolution. However, mechanistic investigations are needed to understand the details of how Hh signaling is transduced in reptiles and the extent to which transcriptional responses are shared between reptiles and more distantly related species.

The brown anole lizard, *Anolis sagrei*, is an emerging squamate model for developmental and functional genetic studies. This species was the first squamate in which *in vivo* gene editing was reported, enabling the production of lizard gene knockouts (Rasys et al., 2019). Additionally, the recent publication of a high-quality reference genome for *A. sagrei* has further enhanced this model system (Geneva et al., 2022). Despite these advances, additional resources and tools are required for mechanistic studies of signaling pathways and cellular processes in *A. sagrei*. Here we characterize the transcriptional responses of *Anolis* and mouse limb bud cells to the Smoothened agonist SAG in cell culture to identify shared and species-specific Hh responsive genes. To enable experimental manipulation of gene function in cell culture, we report the establishment and characterization of a new immortalized *A. sagrei* embryonic fibroblast cell line. As proof-of-principle, we use this cell line to test the role of primary cilia in reptilian Hh signaling by using CRISPR to disrupt the *ift88* gene, which is required for the generation of primary cilia in diverse organisms (Ben et al., 2011; Han et al.,2003; Huang & Schier, 2009; Huangfu et al., 2003; Pazour et al., 2000, 2002). Our results indicate that *ift88* in lizards is required for ciliogenesis and that cilia play important roles in both activation and repression of Hh signaling.

## Results

### Pharmacological induction of Hh signaling in ovo

Previous work has demonstrated that a single injection of Smoothened Agonist (SAG) into pregnant mice at E9.25 or E10.5 is sufficient to activate Hh signaling in the limb buds and induce polydactyly (Fish et al., 2017; Shin et al., 2019). To test our ability to pharmacologically manipulate Hh signaling in anoles, we exposed *Anolis* eggs to a single bolus of SAG within 24 hours of oviposition (*see Methods*). Although it is not feasible to perform timed matings in anoles, *Anolis* embryos within these 24-hour eggs are primarily at Sanger stages 3-4, which are at early stages of limb bud outgrowth and are roughly equivalent to E9.5-10 mouse embryos (Sanger et al., 2008). However, we note that embryos in 24-hour eggs will occasionally be at earlier or later stages of development. To assess whether SAG exposure can induce polydactyly in anoles, SAG exposed eggs were incubated for 15 days to allow digits to develop. Polydactylous embryos were observed in 8 out of 9 eggs treated with 100 µM SAG, with four embryos exhibiting polydactyly in forelimbs and hindlimbs and four embryos displaying polydactyly in only the hindlimbs (Fig. 1B, Table S1). The single SAG treated embryo which did not show polydactyly was at a later stage of development (stage 16 vs. stages 11-14 for the polydactylous embryos), suggesting that the embryo was likely beyond stage 3-4 at the time of SAG exposure. None of the control embryos treated with vehicle (DMSO) alone were polydactylous. Our results demonstrate that, as in mice, SAG can induce polydactyly in anoles.

**Figure 1.**
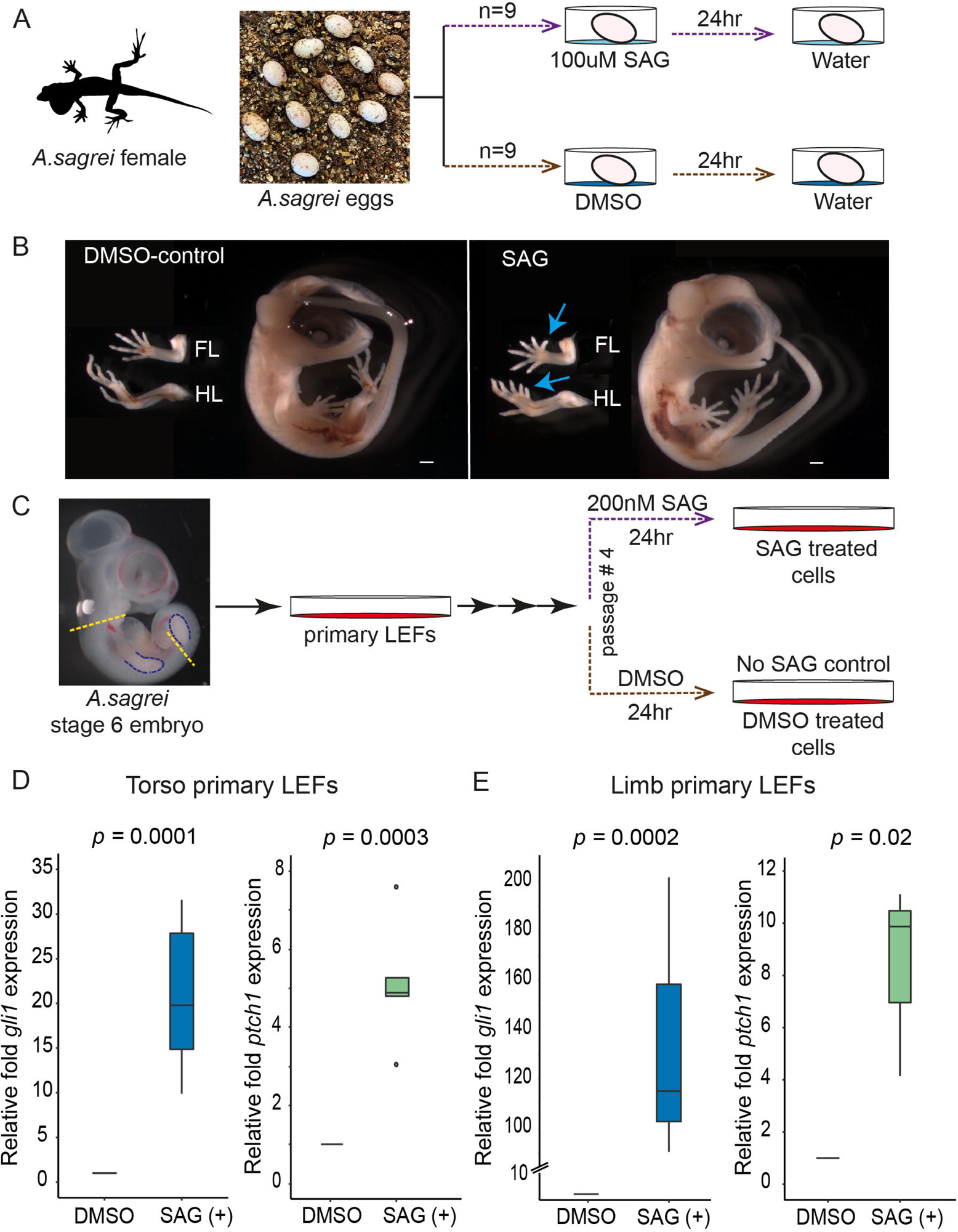
Manipulation of Hh signaling in *A. sagrei* embryos and primary cells. (A) Schematic of *in ovo* treatment of lizard embryos with SAG. Each egg was collected within 24 hrs of being laid and placed on top of a SAG or DMSO solution. These solutions were replaced by water after 24 hrs. (B) Embryos were collected from SAG or DMSO treated eggs after 14 days of incubation. Blue arrows highlight the location of extra digits. FL= forelimbs, HL= hindlimbs. Scale bar = 500 µm. (C) Schematic of primary lizard embryonic fibroblast cell (LEF) SAG treatment. Cells from either the torso (region between dashed yellow lines) or limb buds (outlined in blue) from a single embryo were grown in culture before being treated with SAG or DMSO. (D-E) qRT-PCR of *gli1* (blue) and *ptch1* (green) in the primary torso (D) or limb (E) LEFs exposed to 200nM SAG or DMSO for 24 hrs. Expression was normalized to *gapdh*.

### Comparison of transcriptional responses in A. sagrei and M. musculus after Hh pathway induction

The *in ovo* induction of polydactyly by SAG is consistent with the activation of Hh signaling. To confirm that SAG induces Hh transcriptional responses in anoles, we isolated lizard embryonic fibroblasts (LEFs) from torsos and limbs of late 5/early stage 6 *Anolis* embryos (Fig.1C). Upon SAG exposure, known conserved transcriptional targets of the Hh pathway, *gli1* and *ptch1*, were reproducibly induced in primary LEFs from the torso region and from limb buds. The average induction of *gli1* and *ptch*1 in torso LEFs was approximately 20-fold and 5-fold, respectively (Fig.1D). The induction of these genes in response to SAG was also observed in limb LEFs, with *gli1* and *ptch1* increasing by 136-fold and 8-fold, respectively (Fig. 1E). Having established that SAG can induce the expression of two well-conserved transcriptional targets of the Hh pathway in *Anolis* cells, we next sought to determine the extent to which Hh induced transcriptional responses are shared between *A. sagrei* and *M. musculus*.

To compare Hh induced transcriptional changes in anoles and mice, we stimulated the Hh signaling pathway in primary limb LEFs and primary limb MEFs (mouse embryonic fibroblasts) with SAG. Based on limb bud development, E11.5 mouse embryos are roughly equivalent to *Anolis* embryos at the late 5 to early 6 Sanger stages (Sanger et al., 2008). Primary LEFs and MEFs were collected from stage-matched embryos to generate parallel datasets from these two distantly related species. Each biological replicate of LEFs or MEFs was collected from all four limbs of a single embryo. Culture conditions for the two species were identical except for the temperature, with LEFs being grown at 29°C and MEFs at 37°C. After exposing cells to 200 nM SAG or vehicle (DMSO) alone for 24 hours, RNA was isolated and sequenced. After performing RNA-seq, differentially expressed genes between SAG and DMSO treated cells were identified (Fig. 2A, 2B). We noted that there was more variation in gene expression across our lizard biological replicates than in our mouse replicates (Bartlett’s test of equal variance, p < 2.2 x 10^-16^). Possible reasons for the greater variability in the *Anolis* data include a greater spread in developmental stages or higher levels of genetic variation in the *Anolis* embryos used; the mouse embryos were collected from a single litter while each *Anolis* embryo came from a separate egg laid by a different female. Because of this variability, we used a less stringent significance threshold when identifying SAG responsive genes in lizard than mouse (adjusted *p*-value <0.05 for lizard vs. <0.01 for mouse). We found that 275 genes were significantly upregulated, and 116 genes were downregulated in response to SAG treatment in *Anolis* cells. In mouse, 1056 genes were upregulated, and 844 genes were downregulated in response to SAG.

**Figure 2.**
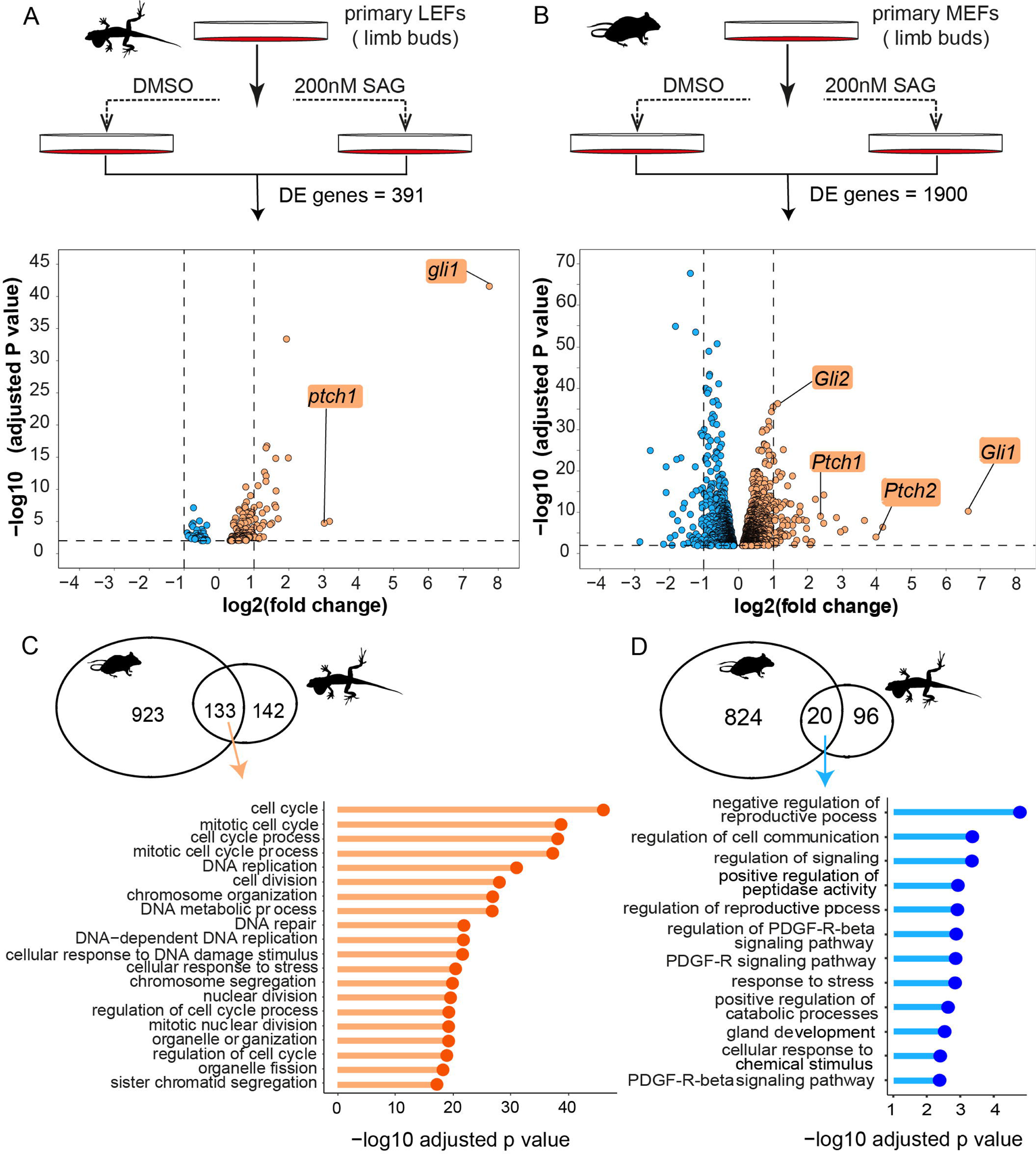
SAG-responsive genes in *Anolis* and mouse primary limb embryonic fibroblast cells. (A-B) Experimental design and volcano plots of differentially expressed (DE) genes in *Anolis* (A) and mouse (B) cells. Volcano plots display DE genes with an adjusted *p* <0.05 for *Anolis* and *p* < 0.01 for mouse. Orange indicates upregulated and blue downregulated in response to SAG. (C-D) Venn diagrams present the number of shared and species-specific upregulated (C) or downregulated (D) genes in *Anolis* and mouse. Lollipop plots show the top biological processes GO terms associated with the shared upregulated or downregulated genes.

A comparison of lizard and mouse SAG responsive genes revealed a total of 181 shared genes (Fig. 2C,2D, Table S2). Of these shared SAG responsive genes, 73% (133 genes) were upregulated in both species and 11% (20 genes) were downregulated in both species. Thus, the majority (84%) of shared SAG responsive genes have a transcriptional response in the same direction in both species. Included among these shared genes are *Gli1*, *Ptch1,* and *Ptch2*, which are all upregulated in LEFs and MEFs in response to SAG and are known direct transcriptional targets of Hh signaling. Similarly, *Hip1* was downregulated in both species upon SAG treatment. Further gene ontology (GO) enrichment analysis of the shared upregulated genes demonstrated that this group is enriched for genes associated with cell cycle regulation (Fig. 2C). A small number of genes responded in opposite directions and are either upregulated in *Anolis* cells yet downregulated in mouse cells (7%) or downregulated in *Anolis* but upregulated in mouse cells (8%). Other genes with different SAG responses include *Gli2,* which is a major activator of the Hh signaling pathway in mice (Bai et al., 2002; Ding et al., 1998; Mo et al., 1997). *Gli2* was upregulated in SAG-treated MEFs but was not responsive in LEFs. Similarly, *Gas1* and *Boc* were downregulated in mouse but did not exhibit a SAG response in *Anolis* cells.

### Generation and characterization of an A. sagrei immortalized cell line for functional genetic studies

We established immortalized *A. sagrei* cell lines to enable functional genetic studies of Hh signaling and other biological processes in reptile cells. To accomplish this, we employed a simple strategy where we serially passaged primary LEFs collected from the torso region of a stage 6 *A. sagrei* embryo (Fig. 3A). Cells entered senescence at passage 26, after which time a small fraction of cells resumed proliferation. Single cells were then sorted into a 96-well plate to generate 40 cell lines from one *Anolis* embryo. To test the responsiveness of these cell lines to SAG manipulations, we chose three cell lines and subjected them to 200 nM SAG treatment for 24 hours. All three clones demonstrated a significant increase in *gli1* and *ptch1* expression as a response to SAG treatment (Fig S1A, B). We chose one of these SAG-responsive cell lines, ASEC-1, to evaluate in greater detail. We observed that *gli1* and *ptch1* displayed a clear trend of elevated expression as SAG concentrations were increased (Fig. 3E, F). When exposed to 200 nM SAG for different time intervals from 6 to 72 hours, the relative *gli1* expression increased with increases in exposure time (Fig S1C, D). In both SAG dosage and time curve experiments, relative *gli1* expression was more sensitive to experimental conditions than *ptch1* expression. We also found that the overall *gli1* and *ptch1* induction values were lower than in primary torso LEFs exposed to similar conditions.

**Figure 3.**
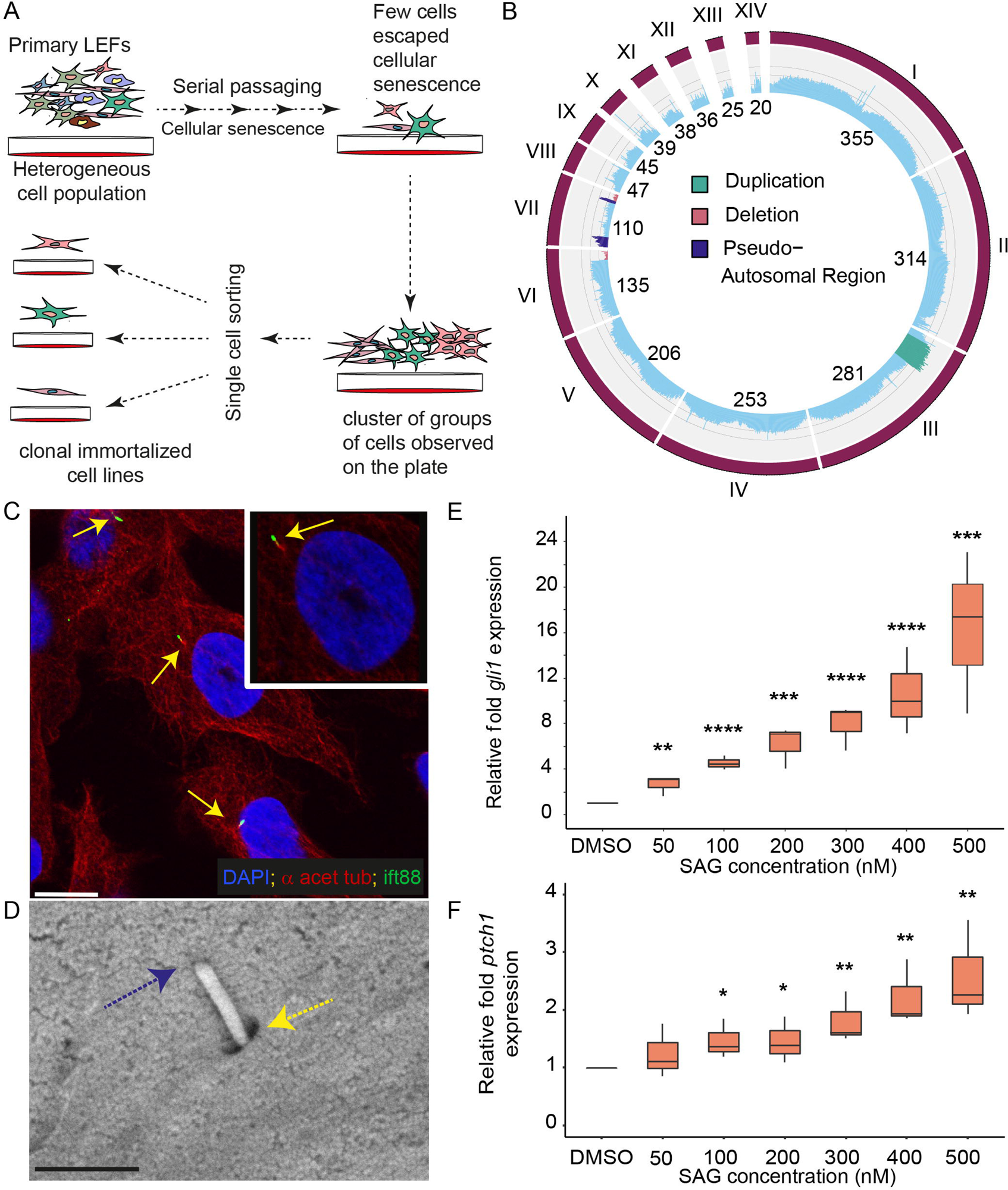
Generation and characterization of the ASEC-1 lizard cell line. (A) Process for the generation of immortalized cell lines from *Anolis* torso primary LEFs. (B) Circular genome plot showing average WGS coverage over 1 MB bins for the ASEC-1 cell line and highlighting potential deletion and duplication events. Grey lines running parallel to axes represent 0.5x, 1x, and 1.5x of the genome wide WGS average. Scaffolds are labeled with Roman numerals. (C) Visualization of the primary cilium on ASEC-1 cells by immunostaining for α acetylated tubulin and ift88. The primary cilium, indicated by yellow arrows, features a ciliary axoneme stained with α acetylated tubulin antibody (red) and the tip of the cilium stained with ift88 antibody (green). Scale bar = 10 µm. (D) SEM image of a primary cilium at 50,000X magnification. A yellow arrow denotes the ciliary pit, and a blue arrow indicates the tip of the cilium. Scale bar = 1 µm. (E-F) qRT-PCR showing relative *gli1* (E) or *ptch1* (F) expression in ASEC-1 cells in response to various SAG concentrations (n=3). Gene expression is relative to DMSO treated cells and is normalized against *tbp* and *atp5f1d*. * *p* <0.05, ** *p* <0.01, *** *p* <0.001, **** *p* <0.0001

To further characterize the ASEC-1 cell line, we assessed the transcriptome (Table S3, S4). RNA-seq analysis detected the presence of transcripts for common fibroblast markers, including *vim*, *col1a1*, *col1a2*, *col5a1*, *lum*, *fap*, *fbln2*, and the cell surface receptor *pdgfra* (Guerrero-Juarez et al., 2019; Meng et al., 2020; Muhl et al., 2020; Sunami et al., 2020). Transcripts for common mural cell markers were also found, namely *des*, *mcam, notch3*, *pdgfrb*, and *anpep* (*cd13*) (Muhl et al., 2020, Sunami et al., 2020). The spindle shaped morphology of the cells combined with the presence of commonly expressed fibroblast molecular markers provides evidence for the fibroblast nature of the cell line (Table 1).

**Table 1:** Fibroblast Marker genes expressed in ASEC-1 cell line identified by transcriptome analysis.

| **Sr.No** | **Gene symbol** | **Base Mean value** |
| --- | --- | --- |
| 1 | *vim* | 373744.26 |
| 2 | *pdgfrb* | 26842.91 |
| 3 | *pdgfra* | 2778.28 |
| 4 | *notch3* | 9239.2 |
| 5 | *mcam* | 15967 |
| 6 | *lum* | 28217.64 |
| 7 | *loxl1* | 1865.38 |
| 8 | *fbln2* | 128.56 |
| 9 | *fbln1* | 527.53 |
| 10 | *fap* | 1021.80 |
| 11 | *des* | 43034 |
| 12 | *dcn* | 56532.66 |
| 13 | *col5a1* | 22997.10 |
| 14 | *col1a2* | 112950.21 |
| 15 | *col1a1* | 69341.19 |
| 16 | *cd34* | 27.83 |
| 17 | *anpep* | 1031.9 |
| 18 | *acta2* | 49194.05 |

To annotate sequence polymorphisms and detect major chromosomal abnormalities in the ASEC-1 cell line, we performed whole-genome shotgun sequencing and aligned the reads to a published reference assembly (Fig. 3B; Geneva et al., 2022). Sequence alignments revealed a large number of polymorphisms that distinguish the ASEC-1 cell line from the *A. sagrei* reference genome (16,101,990 SNPs and 862,358 indels). The annotation of these polymorphic sites provides an important resource for the design of gene editing experiments in ASEC-1. The published reference assembly (AnoSag2.1) was generated from a female and does not contain the *A. sagrei* Y chromosome. However, we found that the genome coverage of the X-specific portion of the X chromosome (i.e., the region of the X that has diverged from the Y chromosome) is half that of the autosomes, suggesting that the cell line is XY and derived from a male embryo. Sex was confirmed using PCR-based genotyping (Fig. S1E). Deviations from the average sequence coverage within a given chromosome indicate likely duplication or deletion events. We noted that scaffold 3 contains an apparent duplication of ∼42 Mb. This region displays 1.5X the average genome-wide coverage. In addition, scaffold 6 exhibits an apparent monoallelic deletion of ∼20 Mb at one end of the chromosome. Scaffold 7, the X chromosome, also contains a putative monoallelic deletion of ∼14 Mb in a pseudoautosomal region. Genes encoding Hh ligands (*shh, ihh, dhh*), Gli proteins (*gli1*, *gli2*, and *gli3*), Hh receptors (*ptch1, ptch2),* and *ift88,* a gene required for ciliogenesis, were not located in duplicated or deleted regions.

Since Hh signaling is functionally linked to the primary cilium in vertebrates (Huangfu et al., 2003; Huangfu & Anderson, 2005), we used immunostaining and SEM to examine ASEC-1 cells for the presence of primary cilia. We co-stained cells with antibodies raised against two established ciliary markers, acetylated α-tubulin and ift88. The staining pattern was consistent with the presence of primary cilia (Robert et al., 2007; Snouffer et al., 2017), with each cell displaying, at most, a single co-stained structure with ift88 staining occurring at the distal tip (Fig. 3C) To confirm the presence of primary cilia, we performed SEM (Fig 3D). SEM images revealed the presence of a ∼1μm long/∼0.3μm diameter protrusion from the cells with a pit at the base, typical of fibroblast primary cilia.

### Targeted disruption of ift88 results in loss of the primary cilium

The *ift88* gene is required for ciliary assembly and maintenance in diverse organisms, ranging from *Chlamydomonas* to mouse (Huangfu & Anderson, 2005; Pazour et al., 2000, 2002). Therefore, to test the requirement of the primary cilium for Hh signaling in anoles, we disrupted the *ift88* gene in the ASEC-1 cell line by CRISPR/Cas9 gene editing. RNA-seq supported gene annotations (Geneva et al., 2022) indicate that the *A. sagrei ift88* gene contains 26 exons with the start codon located in exon 2 (Fig. 4A). We targeted exon 4 to disrupt the ift88 ORF near the N-terminus. After transfecting ASEC-1 cells with *Cas9* and *ift88* gRNA expression constructs, we sorted individual cells into a 96-well plate, which resulted in the isolation of 46 clonal cell lines. PCR amplification of exon 4 followed by PAGE revealed 19 clonal cell lines with indels in *ift88.* Sanger sequencing demonstrated that 17 of these clones carried biallelic frame-shifting mutations (Table S5). In contrast, the chromatograms of two clones (#12 and #35) that appeared wild type by PAGE analysis displayed no mutations. Additional Sanger sequencing of PCR-amplified *ift88* cDNA from mutant clones #28 and #30 confirmed the presence of biallelic frame-shifting mutations in *ift88* transcripts in these clones (Fig. 4B).

**Figure 4.**
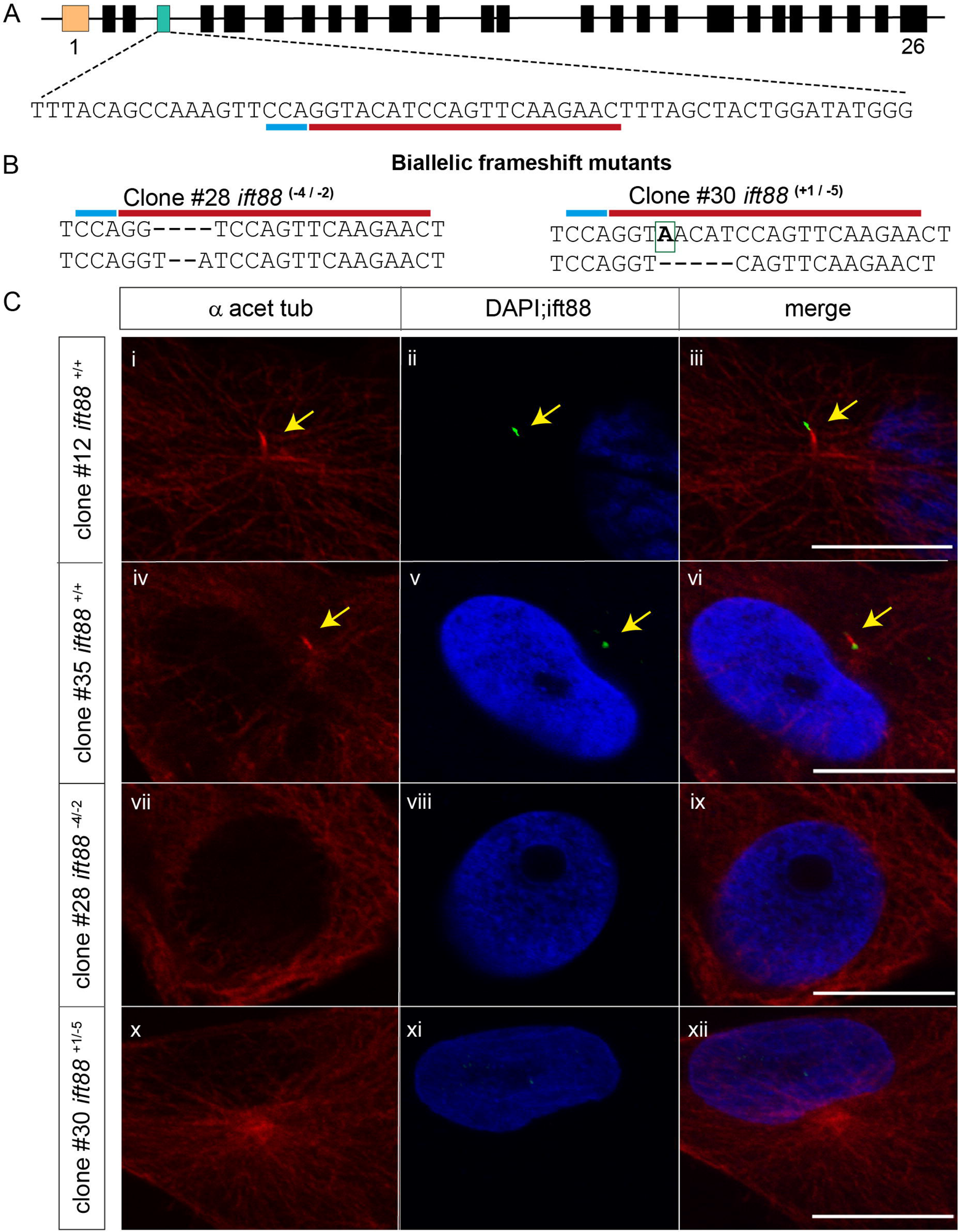
CRISPR/Cas9 editing of the *ift88* gene in ASEC-1 cells. (A) Schematic of the *A. sagrei ift88* gene (not to scale). Rectangles indicate exons. The first exon is noncoding and is indicated in orange. The fourth exon (green) was selected for gene editing. The targeted sequence is underlined in red with the PAM site underlined in blue. (B) Sequence analysis revealed biallelic indel mutations in clones #28 and #30. (C) Representative immunofluorescence images for detection of the primary cilium in wild-type and *ift88* knockout clones. In wild-type clones, α acetylated tubulin (red) stained the primary cilia axoneme with the ift88 (green) staining the tip of the cilium (i-vi). No obvious primary ciliary structures were observed in *Ift88* knockout clones (vii-xii). Yellow arrows indicate primary cilia. Scale bar = 10 µm.

The loss of *ift88* function in ASEC-1 cells is predicted to prevent ciliogenesis. By staining for α acetylated tubulin, we could visualize the primary cilium in at least 70% of cells from the parental ASEC-1 cell line (Fig. 3C, refer methods). Similarly, we could readily identify the presence of primary cilia in two control *ift88^+/+^* ASEC-1 clones (#12 and #35) which did not carry mutations in *ift88* (Fig. 4C). Co-staining with ift88 demonstrated that ift88 is localized at the tip of the cilia. In contrast cells from mutant clones (clone #28: *ift88^-4/-2^*; clone #30: *ift88^+1/-5^*) did not show any ift88 staining. Moreover, based on α acetylated tubulin, roughly 90% of the cells in clones #28 and #30 failed to show ciliary structures (Fig. 4C vii-xii). Approximately 11% of cells in clone #28 and 5% of cells in clone #30 showed ambiguous structures, which could not be confidently categorized as primary cilia.

### ift88 is required for SAG transcriptional responses

To determine whether the loss of *ift*88 and primary cilia affects Hh signaling, we studied SAG responsiveness of *ift88* mutant cell lines. We exposed the parental cell line ASEC-1, clone #12 *ift88^+/+^*, clone#35 *ift88^+/+^*, clone #28 *ift88^-4/-2^*, and clone #30 *ift88^+1/-^*^5^ to 400 nM SAG for 24 hours. For each cell line, vehicle (DMSO) alone was used as a control treatment. The levels of *gli1* and *ptch1* transcripts were quantified relative to the control through qRT-PCR. Both of the WT clones had an increase in *gli1* expression in response to the SAG treatment. However, the increase in relative *gli1* expression was lower than that in the parental cell line (Fig. 5A). The relative levels of *ptch1* expression increased in WT clones, similar to the ASEC-1 cell line (Fig. 5B). Conversely, *ift88* mutant clones #28 and #30 showed no change in *gli1* and *ptch1* expression in response to the SAG treatment (Fig. 5A, B).

**Figure 5.**
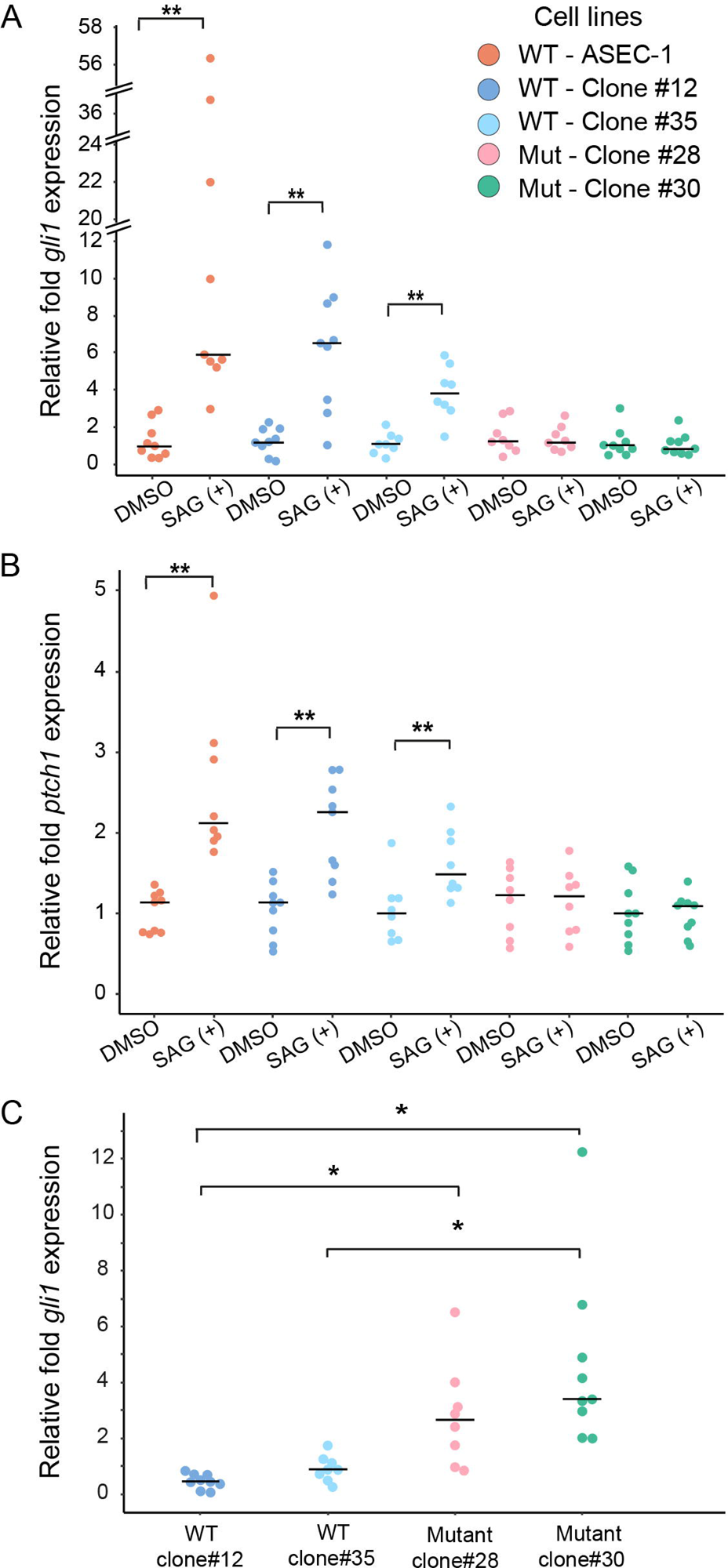
Expression of *gli*1 and *ptch*1 in wild-type and *ift88* knockout ASEC-1 cells. (A-C) Transcript levels were assessed by qRT-PCR. In each cell line, the induction of *gli1* (A) or *ptch1* (B) following SAG treatment was assessed relative to DMSO controls. The parental ASEC-1 cell line as well as two wild-type clones show an increase in *gli1* and *ptch1* expression in response to treatment with 400 nM SAG for 24hrs. Mutant clones #28 and #30 showed no change in *gli1* or *ptch1* expression. (C) Relative basal *gli1* expression in DMSO treated clones. Basal *gli1* expression is relative to the average delta Ct value of *gli1* expression in ASEC-1 DMSO treated cells. *gli1* and *ptch1* expression was normalized to the two reference genes *tbp* and *atp5f1d*. n=9 for each condition. \**p*<0.05, ** *p*<0.01

In mice and zebrafish, the primary cilium also plays a role in repressing the Hh signaling pathway in the absence of ligand stimulation, leading to a higher level of unstimulated, basal Hh pathway activity in mutants lacking cilia (Huang & Schier, 2009; Ocbina et al., 2009). To determine if there is a difference in basal *gli1* expression levels between ASEC-1, WT clones, and *ift88* mutants, we analyzed *gli1* expression in these cell lines relative to the parental cell line in the absence of SAG. We found that there was no statistically significant difference observed between basal *gli1* expression in WT clones #12 and #35 relative to the parental cell line. However, both mutant clones showed elevated basal expression of *gli1* (Fig 5C). Together, our data indicate that *ift88* mutant clones fail to induce the expression of Hh target genes in response to SAG and suggest that cilia are also required in anoles for repression of Hh pathway activity in the absence of inducer.

## Discussion

There are over 11,000 recognized species of squamate reptiles (Uetz P et al.,2023). These species display tremendous diversity in a host of anatomical traits such as limb length, digit morphology, degree of body elongation, and pigmentation (Uetz P et al.,2023). However, mechanistic studies that use functional genetic tools to understand developmental and cellular processes in this species-rich group are lacking. A major barrier has been the absence of effective, inexpensive embryo and gene manipulation tools, methods, and reagents. Encouragingly, *in vivo* gene editing methods have recently been established in three different squamate models (Abe et al., 2023; Rasys et al., 2019; Tzika et al., 2023). Nevertheless, the field of squamate developmental genetics is still new and remains underdeveloped. The aim of the current study was to develop new resources for *A. sagrei* to further advance this species as a reptilian model system for comparative evolutionary, developmental, and functional genetic studies. Specifically, we developed *in vitro* and *in ovo* methodologies in *A. sagrei* to study Hh signaling, which is essential for many developmental processes. Functional analysis of Hh signaling in squamates not only provides an opportunity to understand the role of Hh in reptilian development and regeneration but also can provide an evolutionary perspective on the mechanisms, regulation, and components of the Hh pathway.

### Developmental and transcriptional responses to Hh activation in lizards and mice

To induce Hh signaling in developing embryos, we optimized a method for the uptake of the pharmaceutical compound SAG into eggs by absorption. Hh signaling is known to regulate digit patterning and identity. For instance, polydactyly occurs in several naturally occurring mutants and KO mouse models as a result of increased activation of Hh signaling in the developing limb buds (Anderson et al., 2012; Hill et al., 2003; Lettice, 2003; Lettice et al., 2012; Tickle et al., 1975; Tickle & Towers, 2017). We have found that SAG exposure reliably induces polydactyly in anoles as observed in mice (Fig 1B; Fish et al., 2017; Shin et al., 2019). The specificity and reproducibility of SAG-induced changes in digit number in anole embryos is consistent with hyperactivation of Hh signaling. We speculate that the variation in the number of digits in the forelimb and hindlimb is likely attributable to variation in embryonic stages in 24-hour laid eggs and biological availability of SAG to the developing embryo. In related work reported by Sanger and colleagues, Hh signaling in anoles was manipulated by soaking eggs in cyclopamine, a Hh signaling inhibitor (Sanger et al., 2021). The robustness, simplicity, and cost-effectiveness of *in ovo* exposure to pharmaceutical agents provides a useful tool for the study of Hh signaling at specific embryonic stages. While the absorption of other bioactive compounds into the egg remains to be tested, this general approach may also provide an effective way to perturb other signaling pathways and complements functional analyses based on the generation of constitutive gene knockouts in anoles.

Hh transcriptional responses have been explored in many different developmental contexts and in different model systems (Goodrich et al., 1996; Ingham et al., 2011; Sigafoos et al., 2021). Through these studies, we know that there are a small set of genes that consistently exhibit transcriptional responses to Hh across different tissues and different animal models. Among this core set of Hh responsive genes are components of the Hh signaling pathway itself, including genes that encode PTCH and GLI proteins (Anderson et al., 2012; Chiang et al., 1996; Ding et al., 1998, 1999; Holtz et al., 2013; Hu & Helms, 1999; Karlstrom et al., 1999; Lei et al., 2004; Matise et al., 1998; Mo et al., 1997; Nüsslein-Volhard & Wieschaus, 1980; Sasaki et al., 1997; Shimeld et al., 2007). However, a large proportion of Hh transcriptional responses are cell type specific. Though it has been demonstrated that some cell type specific Hh responses are conserved across species (Bai et al., 2004; Briscoe & Small, 2015; Kicheva et al., 2014; Krauss et al., 1993; Patten et al., 2003; Persson et al., 2002; Ren et al., 2020; Ribes et al., 2010; Stamataki et al., 2005), comprehensive comparisons of Hh induced transcriptional changes in homologous cell populations from different species are largely absent from the literature. Here we performed parallel investigations of Hh induced transcriptional changes in mice and lizards which the tissue source (limb buds), developmental stage, culture conditions, and treatment regimen were carefully controlled. *Anolis* lizards and mice diverged from a common ancestor more than 300 million years ago (Hedges et al., 2006), but share similar pentadactyl digit patterns. Moreover, in both species hyperactivation of Hh signaling during limb bud stages induces polydactyly. Therefore, limbs are ideal for investigating how Hh induced transcriptional changes have evolved across time.

We determined that the mouse orthologs of nearly half of Hh responsive lizard genes also exhibit Hh induced transcriptional changes. Furthermore, the majority of these shared Hh responsive genes (84%) have transcriptional shifts in the same direction in the two species. Thus, our comparison of Hh responsive genes in limb cells from these species has allowed us to reveal the degree to which Hh signaling induces conserved transcriptional responses. Species-specific differences in Hh transcriptional responses may be equally important in understanding how the pathway regulates animal development. The Hh pathway induces transcriptional responses through GLI transcription factors. Upon induction of Hh signaling GLI proteins can be activated through post translational processing as well as through transcriptional upregulation of *Gli* genes. These two mechanisms act to shift the balance of Gli^Act^ to Gli^Rep^ (Falkenstein & Vokes, 2014). It is evident from studies in zebrafish and mice, that the relative importance of the different *Gli* genes can evolve, and the details of how the ratio of Gli^Act^ to Gli^Rep^ is controlled can differ between species or between cell types in an organism (Bai et al., 2002; Huang & Schier, 2009; Karlstrom et al., 2003; Matise et al., 1998; Mo et al., 1997; H. L. Park et al., 2000; Tyurina et al., 2005). While *gli1* transcription is strongly induced by SAG in both mouse and lizard limb cells, we found that *gli2* is induced by SAG only in mice. These results suggest that *gli2* transcriptional regulation by Hh signaling differs between lizards and mice. However, whether the relative roles of the different *gli* genes differ between mice and lizards will require additional functional studies in lizards.

### Establishment of a new cell line for studies of reptilian gene function

To explore developmental mechanisms, gene function, and gene regulation in squamates, a reptilian model system with effective gene manipulation tools is essential. To complement *in vivo* gene editing methods to knockout genes in *A. sagrei* (Garcia-Elfring et al., 2023; Rasys et al., 2019), we developed a resource for functional genetic experiments in cultured fibroblasts to increase the versatility of *A. sagrei* as a reptilian model system. The use of cell culture is ideal for studies that require fine control of the cellular environment and for studies investigating processes occurring at the cellular level (e.g., organelle biogenesis). Cell culture also provides the opportunity to perform high throughput screens of gene function and is ideal for pharmaceutical manipulation of cells.

While the ability to culture and manipulate primary cells is important, immortalized lines provide the ability to perform well controlled studies on genetically homogeneous cell populations. We created an immortalized fibroblast cell line, ASEC-1, by serially passaging *A. sagrei* torso LEFs and isolating clones that escaped senescence (Fig. 3A). Our results demonstrate that ASEC-1 cells can induce Hh signaling in response to SAG and that CRISPR/Cas9 mediated methods can be effectively used to perform gene editing in this cell line. The ASEC-1 transcriptome datasets that we generated provide a deeper characterization of the cell line and are a valuable resource for other researchers who wish to use this line. As is typical for immortalized cell lines, ASEC-1 carries a small number of large duplications and deletions (Rebuzzini et al., 2016). We have defined these chromosomal anomalies to allow researchers to account for these changes when planning experiments. An important additional consideration when using ASEC-1 is that the cell line is derived from wild caught animals from an invasive population in Florida. Since invasive populations of *A. sagrei* have high levels of genetic variation (Bock et al., 2021), we performed WGS on ASEC-1 to annotate the set of polymorphisms present within the cell line. When selecting CRISPR target sites or designing primers, awareness of these polymorphisms is important, and our annotations will facilitate future gene editing experiments in ASEC-1.

### Understanding the role of primary cilia in reptile Hh signal transduction

The *ift88* gene is deeply conserved and is essential for the formation of cilia in species ranging from *Chlamydomonas* to mice (Huangfu et al., 2003; Pazour et al., 2000). In addition, primary cilia are known to be essential for Hh signal transduction in diverse vertebrates, including mouse, chick, and zebrafish (Ben et al., 2011; Huang & Schier, 2009; Huangfu et al., 2003; Huangfu & Anderson, 2005; T. J. Park et al., 2006; Yin et al., 2009) Our demonstration that disruption of *ift88* in lizard cells results in the loss of primary cilia and our conclusion that primary cilia are required for Hh signaling in lizards further support the broad conservation of these roles across vertebrates.

While this study serves as proof of principle that the ASEC-1 cell line can be used to test the role of lizard genes in cellular processes, it also provides the first insights into Hh signal transduction in reptiles. Mutant mouse embryos lacking cilia show a pronounced loss of ligand induced Hh pathway activation. In these mutants, cilia dependent Gli2 activation is lost which in turn dramatically suppresses *Gli1* expression (Haycraft et al., 2005; Huangfu et al., 2003; Huangfu & Anderson, 2005; Jia et al., 2009; Liu et al., 2005). The formation of Gli3 repressor is also greatly reduced in such mutants. These mutants show some mild, ligand-independent (derepressed) activity in the neural tube due to reduced Gli3 repressor (Huangfu et al. 2003). Chicken Talpid3 mutants also lack cilia (Lewis et al., 1999). Such mutants show dramatic loss of expression of Hh target genes in the neural tube, accompanied by ectopic ligand-independent expression of some Hh targets in the limbs. The derepression of some targets is likely explained by reduced ability to generate GLI3 repressor (Davey et al., 2006). In contrast, zebrafish lacking cilia appear to show stronger ligand independent Hh pathway derepression, which cannot be further increased by Hh ligands (Huang & Schier, 2009). The derepression of the pathway in zebrafish *ift88/oval* mutants seems to be mediated by Hh-independent activation of Gli1 protein, rather than by a reduction in Gli repressor activity. We observed a 3-to-4-fold increase in basal *gli1* expression in unstimulated *ift88* mutant lizard embryonic fibroblasts relative to wild-type (Fig 5C), whereas in mouse *Ift88* mutant embryonic fibroblasts (without stimulation) derepression of the Hh pathway was not observed (Ocbina et al., 2009). Thus, it appears that in squamates, similar to ray-finned fish and birds, cilia may play a more important role in ligand-independent repression of the Hh pathway relative to mammals. We speculate that cilia in the shared vertebrate ancestor played an important role in Hh pathway repression, as well as in ligand-induced activation. However, the repressive role of cilia may have become less important in the mammalian lineage after its divergence from reptiles.

Although we have used ASEC-1 to explore Hh signaling here, this cell line can be used to broadly explore gene function in anoles. We note that Zhang and colleagues were able to isolate a subclone of ASEC-1 that can be induced to initiate myogenesis in culture (Zhang et al., 2022). They subsequently used this subclone to investigate the function of the *mymk* gene in lizard myoblasts. As lizards are poikilothermic, the immortalized *A. sagrei* cell lines grow at lower temperatures than mammalian cell lines (29°C vs. 37°C). Therefore, these lines also allow for genetic investigation of homeostatic regulation of cellular processes at lower temperatures. This could be particularly useful for researchers studying the biology of thermoregulation. Moreover, since ASEC-1 was derived from a male (XY) this line may prove useful in addressing questions related to the biology of *Anolis* sex chromosomes and X chromosome dosage compensation mechanisms. For investigators who wish to perform comparative studies of gene function in reptiles, this cell line provides an easily accessible, inexpensive, and renewable resource.

## Materials and Methods

### Animals

*Anolis sagrei* captured in Orlando, FL were housed at the University of Georgia following published guidelines (Sanger et al., 2008). Breeding cages housed up to 5 females and 1 male together, and nest boxes were checked daily to collect eggs within 24 hours of being laid. Outbred ICR background (Envigo) mice were used for collecting mouse embryonic fibroblasts for RNA-seq experiments. All experiments followed the National Research Council’s Guide for the Care and Use of Laboratory Animals and were performed with the approval and oversight of the University of Georgia Institutional Animal Care and Use Committee.

### Induction of Hh signaling via *in ovo* SAG treatment

A total of 18 eggs were collected within 24 hours of being laid. The eggs were carefully cleaned with Kimwipes to remove any debris. In a 24-well plate, each egg was carefully placed in an individual well. A 10 mM SAG stock solution in 100% DMSO solvent was prepared (CAS 912545-86-9 - Cayman Chemical). A 100 µM SAG solution from 10 mM SAG stock was prepared using water as solvent. A control solution with an equivalent volume of DMSO was prepared. The final concentration of DMSO in the control solution was 1%. In each well, 80 µl of either 100 µM of SAG or control DMSO solution was placed below the eggs carefully, ensuring the egg was in contact with the liquid (Fig.1A). 9 eggs received 100 µM of SAG solution and 9 received control DMSO solution. Eggs were incubated at 29°C. After 24 hours, SAG and DMSO solutions were replaced by 80 µl of water. The eggs were incubated for 14 days at 29°C. The eggs were monitored daily and 80 µl of water was added as and when required to avoid drying the eggs. On the 15th day, eggs were dissected in 1X PBS solution, and embryos were examined for morphological differences. Developmental staging was performed according to Sanger et al., 2008. Embryos were imaged and fixed in 4% PFA before being dehydrated in a MeOH series, starting from 25% and ending in 100%. Embryos were stored in 100% MeOH at 4°C.

### Culturing of primary lizard embryonic fibroblasts

*A. sagrei* eggs were cleaned with Kimwipes soaked in 70% EtOH. All debris on the eggshell was removed, and eggs were given 2-3 quick washes with 1X PBS to clean the eggshell. The eggs were dissected in sterile 1X PBS. Individual stage 6 embryos were transferred into a clean petri dish with sterile 1X PBS containing 1X Penicillin-Streptomycin-Amphotericin B solution. The embryo was gently washed by swirling the dish. The process was repeated at least 3 times. For the isolation of LEFs from limb tissue, using sterile forceps, limb buds were separated from the embryo. All four limb buds were aseptically transferred in a 1.5 ml microcentrifuge tube with 0.5 ml 0.05% Trypsin-EDTA. For collecting LEFs from the torso region (Fig 1C), the head, tail, and limb buds were removed with sterile forceps. The rest of the tissue was eviscerated. The tissue was then teased with the forceps and aseptically transferred into a 1.5 ml microcentrifuge tube with 0.5 ml 0.05% Trypsin-EDTA. The tissue was then incubated at 29°C for 45-60 minutes. To facilitate tissue disaggregation, the tissue was teased by pipetting the trypsin solution up and down gently at 15 minutes intervals. After the incubation, the trypsin was neutralized by adding an equivalent volume of LEF growth medium (1X DMEM with 4.5 g/L glucose, without L-glutamine, 10% heat-inactivated fetal bovine serum (Benchmark), 1X glutamine, 1X Penicillin/Streptomycin/Amphotericin B). Cells were collected by centrifuging at 1200-1300 rpm at RT for 5 minutes. Cell pellets were resuspended in 100 µl of LEF growth medium before being uniformly plated in a single well of a 24-well plate and incubated at 29°C at 5% CO_2_ for 1-1.5 hours. This ensured that the cells were attached to the plate. Afterward, 1ml of LEF growth medium was added to each well, and cells were allowed to grow till they became confluent which required around 48-72 hours. This stage was designated passage 0. When the cells became confluent, they were split in a 1:2 ratio and plated again. This was passage 1. Cells from an individual embryo were considered as one biological replicate.

### Pharmacological manipulation of Hh signaling pathway in primary LEFs

Primary limb and torso LEFs were collected from individual *A. sagrei* stage 6 embryos. Primary LEFs from an individual embryo were grown until passage 3. After passage 3, the cells were split and seeded at equal density into two wells of a 24 well plate (passage 4). After the cells became confluent, they were serum starved for 48 hours by adding serum starvation medium (1X DMEM with 4.5g/l glucose, without L-glutamine, 1% heat-inactivated fetal bovine serum (Benchmark), 1X glutamine, 1X Penicillin/Streptomycin/Amphotericin B). Serum starvation results in cell cycle exit inducing ciliogenesis (Pugacheva et al., 2007). 200nM SAG (prepared in serum starvation medium) was added to one well and a DMSO solution (prepared in serum starvation medium) was added to the second well of each biological replicate. 24 hours after the addition of SAG, RNA was collected.

### RNA-seq analysis for DE genes in *A. sagrei* and *M. musculus* after Hh pathway induction

Five *A. sagrei* stage at late 5/early 6 (Sanger et al., 2008) were collected and limb buds were dissected out. Primary limb LEFs were collected as described above. Five embryos from *M. musculus* were collected at stage E11.5. Primary limb MEFs were collected in a similar fashion to the primary LEFs. MEFs were grown at 37°C at 5% CO_2_. For both, LEFs and MEFs from individual embryos were divided into two wells. 200 nM SAG was added following serum starvation for 48 hours to one well and the volume equivalent DMSO solution was added to the second well. The embryos were processed in a pair-wise manner and analyzed as paired sets. We generated 5 biological replicates for each treatment: primary LEFs treated with SAG, primary LEFs treated with DMSO control, primary MEFs treated with SAG, and primary MEFs treated with DMSO control. We collected total RNA 24 hours after the SAG exposure using the mirVana miRNA Isolation Kit (ThermoFisher Scientific). RNA-seq libraries were prepared using the TruSeq Stranded mRNA Library Prep Kit (Illumina) and were sequenced at the Georgia Genomics and Bioinformatics Core. After checking read quality with FastQC v0.11.8, *A. sagrei* reads were aligned to the AnoSag2.1 genome reference (Geneva et al., 2022) and *M. musculus* reads were aligned to the mm10 genome.using HISAT2 v2.1.0. Transcripts were counted using the feature count function from Rsubread v2.10.5 (Kim et al., 2019; Liao et al., 2019). Sample similarity and batch effects by principal component analysis. Bartlett’s test for equal variance was applied to statistically test for differences between gene expression variation in data collected from the two species. Differentially expressed genes between SAG-treated and DMSO-treated samples in both species were identified using DESeq2 v1.36.0. (Love et al., 2014). Batch effects due to sampling from different embryos were corrected by using biological replicates as the batch variable in the design when performing the tests. The RNA-seq data was validated by qRT-PCR in primary LEFs with three biological replicates (Table S6).

### Generation of immortalized lizard embryonic fibroblast cell lines

Primary LEFs from the torso region of stage 6 *A. sagrei* embryo were collected as described above. We adapted the method used for immortalization of MEFs (Xu, 2005). Cells were initially grown in a 24-well plate before transitioning to 10 cm plates, keeping track of the passage number. After the cells were confluent in a 10 cm plate (around passage #6), we serially passaged the cells with a 1:3 split ratio until passage 24. After the 24th passage, we changed the split ratio to 1:6. The growth rate of the cells decreased at this time point as cells started undergoing senescence. At passage 26 very few cells were attached to the plate, and there were no obvious signs of cell growth for 13-15 days. During this time, we frequently changed the LEF growth medium. After this ‘no growth period’, visible cell growth was observed in the form of patches or colonies of cells. The few cells which escaped cellular senescence started growing at this point. We allowed these cells to grow and become confluent. Within 7-8 days the cells in the 10 cm plate were confluent. Cells were then trypsinized and single cells were sorted using the MoFlo Astrios EQ (Beckman Coulter) cell sorter at the Cytometry Shared Resource Laboratory (CTEGD) at UGA. Each cell was sorted into a well pre-filled with LEF growth medium in a 96-well plate. The cells were incubated at 29°C at 5% CO_2_. These single-cell clones started growing in ∼7 days and in ∼15 days became confluent. Once the cells started growing, over a period of 30 days, we expanded these single cell clones from a 96 well plate to a 10 cm plate. We then froze and stored individual lines in liquid nitrogen.

### Immunostaining and detection of primary cilia

In preparation for immunostaining, lizard cells were plated onto a gelatin-coated coverslip in a 24-well plate. Once the cells were confluent, they were serum-starved for 48 hours to promote ciliogenesis, as previously described. After 48 hours, the cells were washed with 1X PBS twice and fixed in EMS grade 4% PFA for 15 min at RT. The cells were then washed with 1X PBS three times. Cells were incubated at RT for 1 hour in freshly made blocking solution (10% heat-inactivated goat serum with 0.1% Triton X100 in 1X PBS). Cells were then washed with wash buffer (1% heat-inactivated goat serum with 0.1% Triton X-100 in 1X PBS) for 10 mins at RT. The cells were incubated with primary antibody solution 1:8000 dilution of acetylated α tubulin mouse monoclonal antibody (Sigma Aldrich T7451) and a 1:125 dilution of ift88 rabbit monoclonal antibody (Abcam ab184566) overnight at 4°C. The following day, cells were washed three times with wash buffer at RT. The cells were then incubated in the dark for 1 hour at RT with anti-mouse cy3 (1:125 dilution, Jackson Immunology 715-167-003), anti-rabbit Alexa 488 fluor (1:125, Invitrogen A21206) and DAPI as a nuclear stain (Sigma Aldrich 1mg/ml). After 1 hour of incubation with the secondary antibodies, we mounted the coverslips on a slide, and VECTASHIELD antifade mounting medium was applied to preserve fluorescence. Slides were stored at 4°C until imaging. The cells were observed and imaged with a Zeiss LSM 880 Confocal Microscope.

Primary cilia were identified based on α acetylated tubulin antibody staining, the position of the nucleus, and the microtubule organization center in context to a ciliary structure. The primary cilia were then confirmed by ift88 antibody staining. A total of 117 cells for ASEC-1, 150 cells for WT clone#12, 137 cells for WT clone#35, 112 cells for mutant clone#28, and 158 cells for mutant clone#30 were analyzed.

### WGS and SNP identification of ASEC cell line

The NEBNext Ultra II FS DNA Library Prep Kit for Illumina was used to prepare libraries for whole genome sequencing of the ASEC-1 cell lines. Libraries were sequenced by Novogene Corporation Inc. Scripts for DNA read alignment are available online at https://github.com/tryggvimcdonald/iLEF-analysis-scripts/tree/main/genome_assembly_scripts. Genome coverage per scaffold was calculated using the pileup.sh command from BBMap (Bushnell, 2014). This same command was also used to calculate the per-bin coverage with various bin sizes. From the coverage of the whole genome, three regions on scaffolds 3, 6, and 7 with deletion or duplication events were identified. The depth command from SAMtools was used to generate average coverage for these regions by dividing the sum of coverage values by the total number of sites. For SNP calling, duplicate reads were first removed using Picard (“Picard Toolkit” 2019). Next, the resultant file was indexed with SAMtools, and a pileup file was created. BCFtools (Li 2011) was used to identify SNPs. SNPs from alignments of low quality were excluded. For detailed methods refer to Geneva et al., 2022. shinyCircos V1.0 (Yu et al., 2018) was used for generation the circular plot.

### CRISPR/Cas9 gene editing of ASEC-1 cell line

An IDT gBlock with a human U6 promoter followed by the *ift88* gRNA sequence (5’-GTTCTTGAACTGGATGTACC-3’) was designed. ASEC-1 cells were transfected with a Cas9 plasmid carrying puromycin resistance (Addgene pSpCas9(BB)-2A-Puro (PX459) V2.0) and the gBlock in a 1:1 ratio using Lipofectamine LTX Plus (ThermoFisher). 48 hours after the transfection, the cells were placed under puromycin selection at a concentration of 25 µg/ml for 24 hours. After 24 hours, puromycin-resistant cells were allowed to grow and become confluent. Cells were then trypsinized, and single cells were sorted into a 96-well plate as described above. The cells were incubated at 29°C at 5% CO2. These single-cell clones started growing in ∼7 days. Once cells started growing, within a time span of 30-40 days we expanded the clones, froze them, and stored them in liquid nitrogen for later analysis.

Cell lines were screened for indel mutations by PCR amplifying the targeted *ift88* region with the primers listed in Table S7. PCR conditions were 95°C 2 min; 30 cycles of 95°C 30sec - 58°C 30 sec - 72°C 30 sec; 72°C – 5 min for final extension. PAGE protocol was adapted from VanLeuven et al. (2018). We performed electrophoresis at 150 mV for 7 hours. Following PAGE screening, Sanger sequencing was performed on the PCR products to precisely identify sequence alterations in each cell line. Multiplex PCR-based sex genotyping of the ASEC-1 cell line was performed using primers listed in Table S7. The PCR conditions were 95°C for 30 sec for initial denaturation; 35 cycles of 95°C for 30 sec; 58°C for 30 sec; and 72°C for 30 sec; 72°C for 2 min for the final extension.

### RNA isolation and qRT-PCR

Cells were lysed in 500 µl Invitrogen TRIzol reagent and total RNA was isolated according to the Invitrogen TRIzol reagent user guide. RNA yield was measured using Qubit™ RNA High Sensitivity assay kit. cDNA was synthesized using ProtoScript® II First Strand cDNA Synthesis Kit (New England BioLabs). *gli1* and *ptch1* mRNA expression in the cells exposed to SAG were measured by qRT-PCR using Roche the LightCycler® 480 instrument. The LightCycler® 480 SYBR Green I Master (Roche) mixed with the respective primers and 10 ng of cDNA template was used for individual reaction. The PCR conditions were as follows: Pre-incubation at 95°C for 5 mins followed by 45 amplification cycles of 95°C for 10 sec, 58°C for 10 sec, and 72°C for 10 sec. For each biological replicate, three technical replicates were used. For primary LEFs delta Ct values were calculated by normalizing *gli1* and/or *ptch1* expression with *gapdh* as the reference gene. For the remaining experiments performed, *tbp and atp5f1d* were used as the reference genes.

RNA-seq dataset was used to identify and select reference genes for qRT-PCR. Transcript levels of *gli1* and *ptch1* were comparable with *tbp* and *atp5f1d*. The RNA-seq data indicates that *tbp* and *atp5f1d* were stably expressed in DMSO and SAG-treated LEF samples. The stable expression was defined as a coefficient of variance below 0.5 within the biological replicates. We then confirmed stable expression of these genes in ASEC-1 cell line by analyzing Ct values between biological replicates when exposed to SAG and DMSO treatments. When more than one reference gene was used as a normalizer, the geometric mean of delta Ct values with the individual normalizer gene was calculated. This geometric mean of delta Ct value was then further used for 2^-delta-delta Ct calculations. Statistical analyses were performed on delta Ct values using paired *t* test (Fig 1, Fig 3,Fig S1), Wilcoxon signed-rank test, the Kruskal-Wallis rank sum test and Kruskal-Wallis multiple comparison test (Fig 5).

### Pharmacological manipulation of the Hh signaling pathway in ASEC-1

A 24 well plate was seeded with an equal density of the ASEC-1 cells. Each well was considered a biological replicate. Cells were prepared and serum starved as described above. Cells were treated with SAG concentrations ranging from 50 nM to 500 nM. Control wells were treated with 0.05% vehicle (DMSO). For every SAG concentration, three biological replicates were assayed, and RNA was collected after 24 hours of SAG treatment. In separate experiments, we exposed the ASEC-1 cell line to 200 nM SAG for time intervals ranging from 6 to 72 hrs. For each time interval, an equivalent volume of vehicle (0.02% DMSO) was used as a control. These experiments were repeated twice. For testing the SAG responsiveness of WT and *ift88* subclones, three biological replicates were treated with 400 nM SAG, and three biological replicates were treated with 0.04% DMSO for 24 hours. This experiment was repeated three times.

### Scanning Electron Microscopy

ASEC-1 cells were plated on a gelatin-coated coverslip in a 24-well plate. Once the cells were confluent, we serum-starved them for 48 hours to promote ciliogenesis, as described previously. After 48 hours, the cells were washed carefully with 1X PBS at least 3 times to remove all the traces of serum. The cells were then fixed in 2% glutaraldehyde and 2% PFA. The cells were processed and imaged by the core facility using the Thermo Fisher Scientific Teneo, a field emission scanning electron microscope.

### Accession number

The accession number for the RNA-seq data and WGS data generated for this work is GEO: GSE242114.https://www.ncbi.nlm.nih.gov/geo/query/acc.cgi?acc=GSE242114

## Acknowledgment

We thank Jacob Burnett and Bidushi Chandra for their initial exploration of *A. sagrei* cells in tissue culture that helped to inspire this work. In addition, we thank Christina Sabin, Anna Iouchmanov, Rida Osman, Sneha K. Mohan, Heike Kroeger, and Aaron Alcala for their comments on this manuscript. The graphic icons of the lizard and mouse were kindly provided by Aaron Alcala. This work was funded by the National Science Foundation award 1827647 to D.B.M and J.E.

## Supplementary Data

**Figure S1. SAG responsiveness of immortalized cell lines assayed by qRT-PCR and sex genotyping of ASEC-1.**

(A-B) Relative fold *gli1* and *ptch1* expression in three immortalized clonal cell lines generated ASEC-1 (n=3), ASEC-4 (n=2), and ASEC-5 (n=3). The bar graph represents *gli1* and *ptch1* induction in SAG treated samples relative to DMSO control. The data was normalized to *gapdh* expression. For each biological replicate, there were three technical replicates. (C-D) Relative fold *gli1* and *ptch1* expression in the ASEC-1 cell line in response to 200 nM SAG exposure for different time intervals (n=6). The box plots represent *gli1* and *ptch1* induction in SAG treated samples relative to DMSO control for that time point. The data was normalized using two reference genes *tbp* and *atp5f1d*. * *p*< 0.05, ** *p*<0.01, *** *p*<0.001, **** *p*<0.0001. *p* values represent the statistical significance between delta Ct values of DMSO and SAG treated samples by paired *t* test (E) PCR genotyping results confirming the presence of Y chromosome in the ASEC-1 cell line.

**Table S1: Digit numbers in polydactylous embryos from the eggs treated with 100 µM SAG**

**Table S2: Differentially expressed genes in response to 200 nM SAG exposure for 24 hr in *A. sagrei* and *M. musculus* primary limb cells**

**Table S3: Differentially expressed genes in response to 400 nM SAG exposure for 48hr in ASEC-1 cell line**

**Table S4: Upregulated Hh responsive genes in ASEC-1 cell line after 400 nM SAG exposure for 48hr.**

**Table S5: Number of *ift88* mutants clones and types of indels**

**Table S6: Validation of RNA-seq data by qRT-PCR**

**Table S7: List of primer sequences used**

## Notes

### Competing Interest Statement

The authors have declared no competing interest.

