## Supplementary figures and images for "A new cell culture resource for investigations of reptilian gene function"

### Supplemental Figure 1

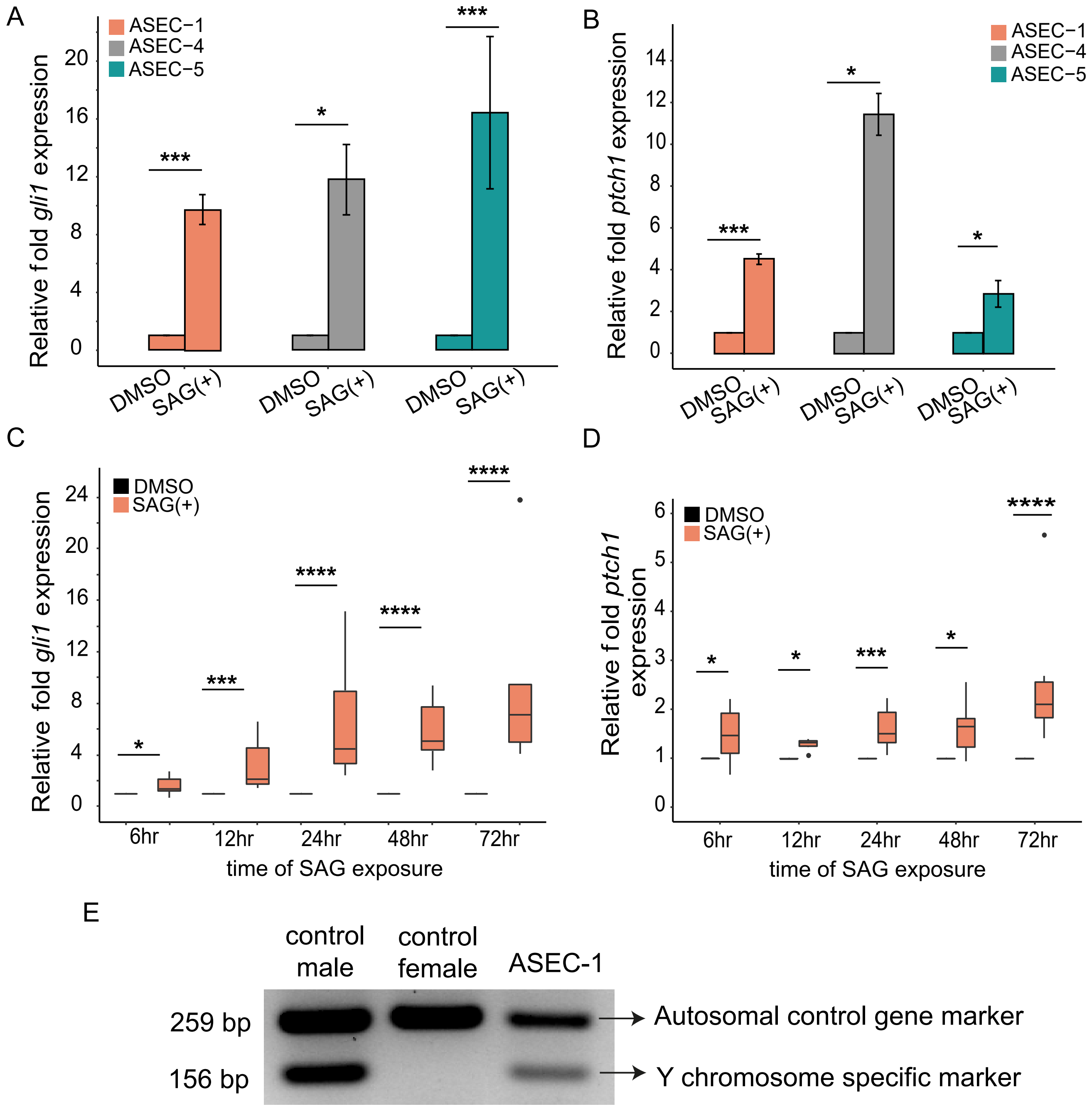
