## Supplemental Table S1 for "A new cell culture resource for investigations of reptilian gene function"

**Table S1: Digit numbers in polydactylous embryos from the eggs treated with 100 µM SAG**

| **No.** | **Number of digits on hindlimbs** | | **Number of digits of forelimbs** | | **Developmental stage** |
| --- | --- | --- | --- | --- | --- |
|  | Left | Right | Left | Right |  |
| 1 | 5 | 5 | 5 | 5 | Late 16 |
| 2 | 6 | 6 | 5 | 5 | 13 |
| 3 | 6 | 6 | 6 | 6 | Late 13 |
| 4 | 6 | 6 | 6* | 5* | 14 |
| 5 | 6 | 6 | 6 | 6 | 13 |
| 6 | 6 | 6 | 5 | 5 | 14 |
| 7 | 6 | 6 | 5 | 5 | 13 |
| 8 | 6* | 9* | 6 | 6 | Late 11 |
| 9 | 6 | 6 | 5 | 5 | Late 13 |

* Asymmetry observed between left and right digit numbers
