## Supplemental Table S4 for "A new cell culture resource for investigations of reptilian gene function"

**Table S4: Upregulated genes in response to 400 nM SAG exposure for 48hr in ASEC-1 cell line.**

| **Gene symbol** | **Protein encoded** | **Fold induction** | **Regulated by Hh signaling in other systems** | **References** |
| --- | --- | --- | --- | --- |
| *gli1* | Gli transcription factor | 4.96 | induced in vertebrates | Lee et al., 1997; Vokes et al., 2007 |
| *vcan* | versican,  ECM-associated proteoglycan | 4.81 | induced in chicken  gut mesenchyme | Nagy et al., 2016 |
| *ndnf* | Neuron-Derived Neurotrophic Factor | 2.14 | induced ndnf-like factor *nord* in *Drosophila* | Yang et al., 2022 |
| *scube3* | Signal peptide-CUB-EGF domain-containing protein 3 | 2.02 | yes, in murine fibroblasts possibly regulated hair follicle growth | Liu et al., 2022 |
| *ptch1* | Hedgehog receptor | 1.79 | induced in vertebrates | Goodrich et al., 1996; Marigo et al., 1996 |
| *bgn* | biglycan,  ECM-associated proteoglycan | 1.72 | induced in colorectal cancer cells, possible regulation | Zeng et al., 2022 |
| *col4a1* | Collagen 4a1, structural component of basement membranes | 1.57 | yes, possible regulation in Shh-medulloblastoma | Northcott et al., 2011 |
| *ptch2* | Hedgehog receptor | 1.56 | induced in vertebrates | Holtz et al., 2013 |
| *vax1* | Developmental transcription factor | 1.39 | induced in vertebrates | Hallonet et al., 1999 |
