## Supplemental Table S5 for "A new cell culture resource for investigations of reptilian gene function"

**Table S5: Number of *ift88* mutant clones and types of indels**

| **No.** | **Type of indel** | **Number of clones** | **Biallelic and out-of-frame mutation** |
| --- | --- | --- | --- |
| 1 | + 1 / - 5 | 5 | yes |
| 2 | - 1 / - 7 | 1 | yes |
| 3 | - 2 / - 4 | 1 | yes |
| 4 | + 1 / - 2 | 1 | yes |
| 5 | + 1 / - 4 | 1 | yes |
| 6 | + 1 / - 1 | 3 | yes |
| 7 | + 1 / - 9 | 6 | No |
| 8 | + 227 / (WT/-1?) | 1 | No |
