## Supplemental Table S6 for "A new cell culture resource for investigations of reptilian gene function"

**Table S6: Validation of RNAseq data by qRT-PCR**

| **Embryo no.** | **Fold induction relative to DMSO control** | | | |
| --- | --- | --- | --- | --- |
|  | *gli1* | *ptch1* | *cldn1* | *ramp2* |
| 1 | 12.69 | 2.81 | 1.37 | 1.64 |
| 2 | 65.79 | 9.94 | 2.16 | 3.81 |
| 3 | 66.02 | 9.56 | 1.25 | 3.52 |
| ***p*-value =** | 0.00014 | 0.00408 | 0.01709 | 0.00729 |

*p* values are associated with a two-tailed paired *t*-test. The analysis was performed using delta ct values.
