## Supplemental Table S7 for "A new cell culture resource for investigations of reptilian gene function"

**Table S7: List of primer sequences used**

| **Gene name** | **Primer name** | **Sequence** |
| --- | --- | --- |
| RT-qPCR primers | | |
| *gapdh* | A.car-gapdh-fwd | ATCGGAGTCAACGGATTTGG |
|  | A.car-gapdh-rev | CATGTAGACCATGTAGTTCAGG |
| *tbp* | A.sag-tbp-fwd | TCTCCAATGACTCCCATGAC |
|  | A.sag-tbp-rev | CAGCCAAGATTTACCGTAGA |
| *atpf1d* | A.sag-atpf1d-fwd | AGACTCTTCCGTCCAACTCC |
|  | A.sag-atpf1d-rev | TCAGACAAGGCCTTCTCCAG |
| *gli1* | A.sag-gli1-fwd | GCTCAGTACATGCTGGTTGTC |
|  | A.sag-gli1-rev | CCGTGAGTAGGCTTTATTGCAG |
| *ptch1* | A.sag-ptch1-fwd | GTGGAGTTTACGGTTCACATTG |
|  | A.sag-ptch1-rev | CACAGGTGCAAACATGTGTTCT |
| *ramp2* | A.sag-ramp2-fwd | AGTGGCAGGGTGAGCAAAG |
|  | A.sag-ramp2-rev | GGTCCTCCTTCCAAAGACAA |
| *cldn1* | A.sag-cldn1-fwd | TGAAGTCAAGAAGATGAGGATG |
|  | A.sag-cldn1-rev | AGTGAATGGGTTGAAGAACTC |
| Genotyping and sequencing primers | | |
| *ift88*  (PAGE primers) | ift88-screen-F1 | GATAATTCTGAATTAATTATAAACTCTTG |
|  | ift88-screen-R1 | TAACATTTAGGGCTCACTGG |
| *ift88*  (For Sanger sequencing) | ift88-screen-F2 | GTTATCAGTAGGAGGCAGTC |
|  | ift88-screen-R3 | CTCACTGGCTGAAATGCTGAG |
| *ift88*  *(*cDNA Sanger sequencing) | *Asag_ift88_Ex1-F1* | CTGAAGCAGATGAAGATGATC |
|  | *Asag_ift88_Ex8-R3* | TCCAGCATCTTTTGCCTTCT |
| *ift88*  (Illumina sequencing) | *AS_ift88_illuminaseq_F2* | ACACTCTTTCCCTACACGACGCTCTTCCGATCTGTTATCAGTAGGAGGCAGTC |
|  | *AS_ift88_illuminaseq_R3* | GACTGGAGTTCAGACGTGTGCTCTTCCGATCTCTCACTGGCTGAAATGCTGAG |
| Sex-genotyping primers | | |
| *AsagB-F1* | | GAAGACCAGGAGAGCAARGTC |
| *AsagB-R1* | | GATGTCGGCAGCYTTGCGTAC |
| *Kanc1-ACF* | | CCTTCCTTTGTAGGATCCAGTG |
| *Kanc1-ACR* | | GGAGCACAGGGATAGTTTTGAC |
